# How sensory load shapes the neural processing and perception of visual durations

**DOI:** 10.64898/2026.05.08.723690

**Authors:** Francesca Iris Bellotti, Marco Zanon, Domenica Bueti

## Abstract

The sensory content and temporal structure of stimuli have been shown to consistently bias duration perception. Temporal intervals filled with continuous sensory input (“filled intervals”), are often perceived as lasting longer than intervals marked only by their onset and offset (“empty intervals”). Despite this robust behavioral finding, it remains unclear whether filled and empty intervals rely on similar or distinct neural mechanisms and, more generally, how sensory format shapes the neural processing of millisecond time. To address this question, we asked twenty-one healthy participants to reproduce visual durations across different stimulus configurations while high-density scalp EEG was recorded. Behavioral results revealed differences in performance across stimulus configurations. Event-related potentials (ERPs) recorded at occipito-parietal and fronto-central electrodes between 0.1 and 0.4 s after duration offset were modulated in amplitude by both stimulus duration and format. These modulations scaled with the sensory load of the stimulus and its duration, suggesting a common underlying mechanism. A Representational Similarity Analysis (RSA) of the ERP data showed that *perceived* time was represented more strongly than physical time particularly at occipito-parietal electrodes, but only within the 0.2–0.3 s post-offset window, where stimulus format exerted a pronounced effect on the ERP signal. These findings highlight the role of sensory processing in shaping duration perception and its neural coding, and reveal an early neural signature of perceived time in occipito-parietal electrodes.

**Significance statement:** Our perception of subsecond durations is distorted by the sensory content of stimuli. Here, we investigated how stimulus configuration shapes the neural correlates of visual duration perception. Specifically, we asked whether temporal intervals filled with continuous sensory input are processed differently from those lacking such content. We found that, between 0.2 and 0.3 s after interval offset, ERP amplitudes were modulated by stimulus content, and in this same temporal window the EEG signal reflected the perceptual bias. These findings support the view that duration processing and perception are deeply rooted in sensory processing.

## 2 Introduction

Distortions of time perception are a common experience in daily life: waiting for the bus feels endless, while a dinner with friends flies by. Even at the sub-second scale, research has shown that perceived time often deviates from physical time (Eagleman, 2008). Psychophysical evidence reveals that the perceived duration of stimuli is biased by various non-temporal features: in the visual domain, for instance, brighter, larger, and faster stimuli are perceived to last longer than their dimmer, smaller, and slower counterparts (Benton & Redfern, 2016; Kaneko & Murakami, 2009; Xuan et al., 2007). This evidence points to the amount of sensory information provided by stimuli as a crucial factor in perceived duration. A specific instance of this idea is the “filled duration illusion” phenomenon, whereby intervals that consist of continuous sensory inputs (“filled intervals”) are perceived as longer compared to intervals only marked by two brief events (“empty intervals”) of equal duration (Horr & Di Luca, 2015; Thomas & Brown, 1974; Wearden et al., 2007). Besides effects on mean perceived duration, filled and empty intervals also differ in discrimination performance (Grondin, 1993) and learning effects (Li et al., 2022). These differences raise the question of whether or not empty and filled intervals rely on the same timing mechanism.

Early accounts of the filled duration illusion, grounded in pacemaker-counter models, attributed these effects to an increased pacemaker rate during filled intervals, resulting in a greater accumulation of temporal “ticks” and therefore a longer perceived duration (Santi et al., 2006; Wearden et al., 2007). However, recent perspectives on temporal processing challenge the notion of a centralized clock. Instead, they emphasize sensory-based mechanisms, where the encoding of duration is inherently tied to the sensory coding of stimuli (Bueti, 2011; Centanino et al., 2026; Reinartz et al., 2021) and to the integration of sensory evidence in time (Ofir & Landau, 2022; Toso et al., 2021). For instance, Toso et al. (2021) demonstrated that distortions in perceived duration are linked to activity in the primary somatosensory cortex (S1) of rats, which encoded the non-temporal feature (vibration intensity) driving the temporal illusion. Similarly, optogenetic manipulations of S1 activity can selectively dilate or compress perceived duration, further supporting the role of sensory cortices in generating temporal percepts (Reinartz et al., 2021). In humans, a study by Tonoyan et al. (2022) investigated the compression in perceived duration induced by motion adaptation and identified its electrophysiological correlates in the amplitude of the N200 component at occipital electrodes and beta-power modulation in the early phase of the stimulus presentation. These results highlight the contribution of early, low-level sensory processing in determining our subjective sense of time. Accordingly, the filled duration illusion may arise from differences in the sensory encoding of stimuli. Despite its robustness, however, the neural underpinnings of the filled duration illusion have received limited attention.

To our knowledge, only one study investigated this phenomenon at the neural level in the sub-second range (Pfeuty et al., 2008). In this study, participants performed an auditory duration discrimination task while electrophysiological activity was recorded and compared between filled and empty stimuli. The results revealed that the Contingent Negative Variation (CNV) component was significantly larger for filled stimuli than for empty stimuli. The authors interpret this finding as evidence that sensory content influences temporal processing. However, the authors also acknowledged that the observed differences in CNV amplitude could reflect the superimposition of temporal processing activity and sensory-sustained activity in the filled condition, rather than a genuine difference in clock rates. Moreover, and most critically, their analysis did not examine whether these neural differences were directly linked to differences in perceived duration. This missing link leaves open the question of whether and how neural activity differences between filled and empty intervals translate into differences in subjective time perception.

Here, we aimed to investigate the neural correlates of the temporal processing for visual stimuli characterized by different levels of sensory load: filled, flanked (i.e. filled intervals with sharp onset and offset) and empty. In addition, we sought to link perceptual distortions measured behaviorally to the EEG signal. To this end, we tested participants in a temporal reproduction task with filled, flanked, and empty stimuli while recording high-density EEG activity.

## 3 Materials and Methods

### 3.1 Participants

Twenty-one healthy participants (15 females, 6 males) took part in the experiment. Participants’ ages ranged from 19 to 35 years (mean: 26.20, SD: 4.74); they were right-handed and had normal or corrected-to-normal visual acuity. All participants provided written informed consent and received financial compensation for their participation. All experimental procedures were approved by the Ethics Committee of the International School for Advanced Studies (SISSA), and were in line with the Declaration of Helsinki (protocol nr. 10035-III/13).

### 3.2 Experimental design

#### Stimuli

Stimuli were generated in MATLAB R2019b using the Psychophysics Toolbox extensions (Kleiner et al., 2007). The stimuli consisted of flickering white noise patches presented within a circular aperture of 4° of visual angle, against a gray background (figure 1). The stimuli were presented with two different levels of contrast. The contrast manipulation was included to examine whether contrast, in addition to stimulus configuration, could elicit perceptual bias as reported in previous works (Benton & Redfern, 2016). The contrast was defined by the separation between two Gaussian distributions from which pixel values were sampled. The boundary values for each contrast condition were computed using a linear transformation, ensuring that pixel intensities were symmetrically distributed around the gray background according to the specified contrast level. In the filled condition, the stimulus was continuously displayed for the entire duration of the interval, such that the time span was occupied by an ongoing sensory event. In this condition, the separation between the Gaussians corresponded to Michelson contrast values of either 15% or 30%. In the flanked condition, two additional high-contrast frames (33,3 ms each) were presented at the beginning and end of the interval, serving as temporal markers. The separation between the Gaussians for these flankers corresponded to Michelson contrast values of either 45% or 90%. In the empty condition, duration was defined by the silent gap between the two flanker frames, without continuous visual input in between them except a fixation cross. The fixation cross covered 0.3° of visual angle. The different stimulus types (filled, flanked and empty) were presented in separate blocks of trials.

**Figure 1:**
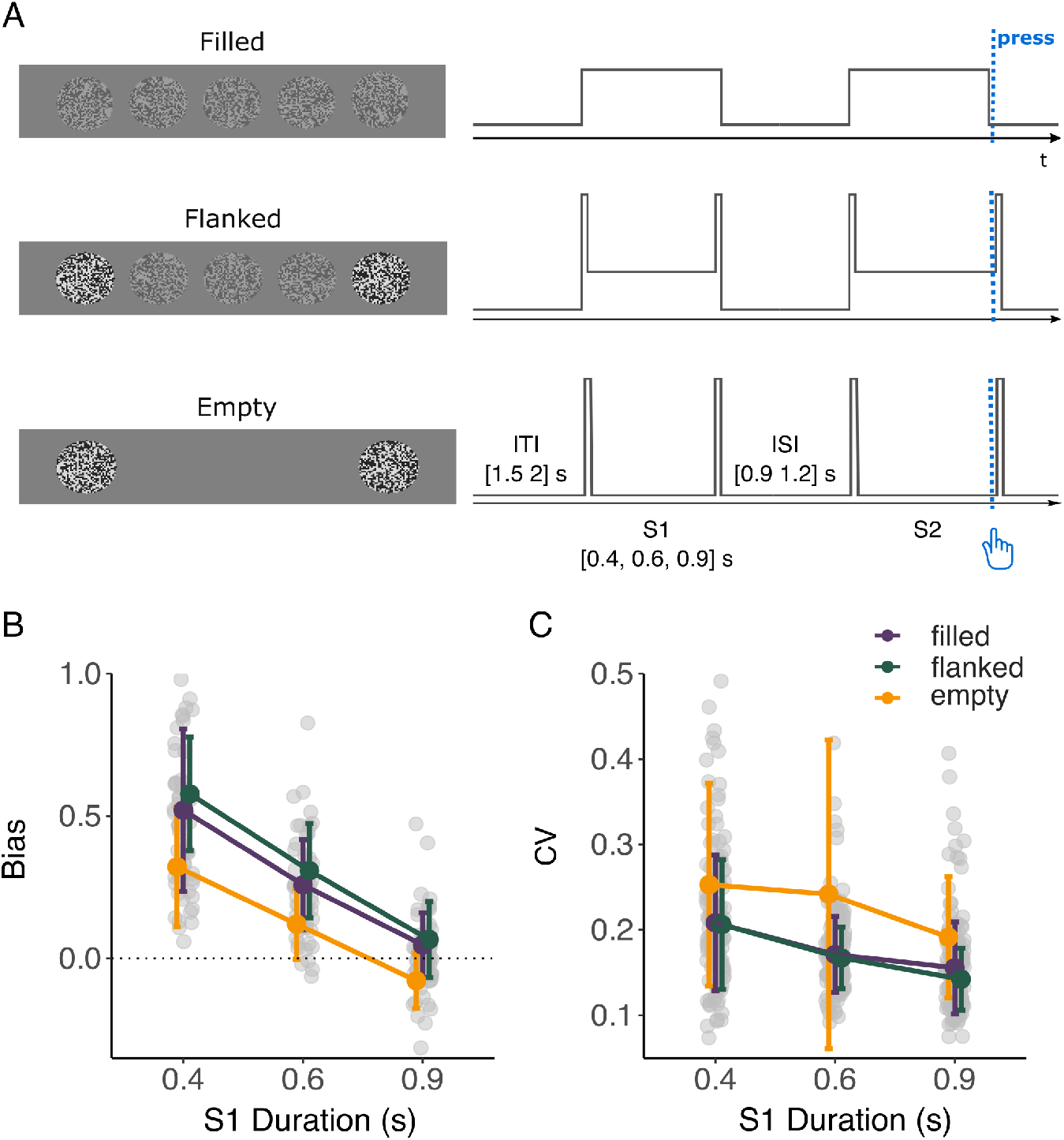
Experimental design and behavioral results.**A)** Frame-by-frame representation of stimulus types and schematic representation of a trial in each stimulus condition. In each trial, a template stimulus (S1) was presented for 0.4, 0.6 or 0.9 s. After a variable interstimulus interval, sampled from a uniform distribution from 0.9 to 1.200 s, a second stimulus appeared. Participants were instructed to press the space bar whenever they thought S1 duration had elapsed. Participants’ press triggered the offset of the reproduction stimulus, i.e. the disappearing of the stimulus in the filled condition, or the appearance of a transient stimulus in the flanked and empty condition. **B)** Mean bias of each participant across S1 durations, divided by stimulus type. Bars indicate the Standard Deviation. A positive bias indicates over-reproduction, whilst a negative bias indicates under-reproduction of duration. **C)** Mean Coefficient of Variation (CV) across S1 durations, divided by stimulus type. Bars indicate the Standard Deviation.

#### Procedure

The experiments were conducted in a quiet and darkened room, using a BenQ XL2720-B monitor. The screen resolution was 1920 × 1080 pixels and its refresh rate 60 Hz. The participants performed the task sitting in front of the screen at a viewing distance of 57 cm and with their head on a chin rest. Participants were tested in a duration reproduction task, schematically represented in Figure 1A. Each trial consisted of the sequential presentation of two stimuli S1 and S2. S1, the first stimulus of the pair, was the actual duration participants had to judge and later reproduce. This stimulus could last for one of 3 distinct durations i.e., 0.4, 0.6 or 0.9 s. This stimulus could be a filled, a flanked or an empty interval. After a variable inter-stimulus interval (ISI), ranging between 0.9 and 1.2 s, a second stimulus appeared (i.e., S2), in the same format (i.e. filled, flanked or empty) as S1. Participants had to press a key on the keyboard to mark the offset of S2 whenever they judged its duration to match that of S1. The inter-trial interval (ITI) ranged from 1.5 to 2 s. Stimulus contrast in S2 was the same in all trials; the filled patches were at 22.5% Michelson’s contrast, while the flanker patches were at 70% Michelson’s contrast. At the beginning of the experimental sessions, participants familiarized with the task for a total of 30 trials (10 trials for each stimulus condition). Only during this training a feedback was provided at the end of each trial. In the experimental session, participants completed three blocks of trials, one for each stimulus type. Each block consisted of 150 trials (50 trials for each S1 duration), for a total of 450 trials. The order of blocks was randomized across participants.

### 3.3 Analysis

#### 3.3.1 Behavioral data analysis

We analyzed participants’ reproduction performance to assess how stimulus configuration affected perceived duration. First, trials in which the reproduced duration exceeded 3 s or fell below 0.1 s were excluded from the dataset. This process resulted in the removal of 0.22% of trials. Two behavioral measures were taken for each participant. The first was a normalized measure of reproduced duration, referred to as ‘bias’, computed as follow: *bias* = (*R*_*d*_ − *T*_*d*_)*/T*_*d*_, where *R*_*d*_ is the reproduced duration and *T*_*d*_ is the S1 duration. A positive value of bias indicates an over-reproduction, while a negative bias indicates an under-reproduction. The second measure was the coefficients of variation (CV), which was computed as follow: *CV* = *R*_*std*_*/R*_*mean*_, where *R*_*mean*_ is the mean reproduced duration across trials of the same condition (for each participant, stimulus type and S1 duration) and *R*_*std*_ is the corresponding standard deviation. To quantify the effects of stimulus type on bias and CV values, we employed linear mixed-effect regression models (LME, R package *lme4*, Bates et al., 2015), incorporating *stimulus type, S1 duration* and *contrast* as fixed predictors and treating *participants* as a random intercept factor.

#### 3.3.2 EEG recordings and pre-processing

While participants performed the task, EEG was continuously recorded from 64 passive electrodes (EasyCap, GmbH, Germany) at 1000 Hz sampling rate with an actiCHamp Plus amplifier (GmbH, Germany). The electrodes were mounted on an elastic cap and placed according to the international 10-20 system. Two additional electrodes were placed for horizontal electro-oculogram (EOG, one electrode at the right eye corner) and for referencing (right mastoid). All sites were grounded to AFz and impedance of all electrodes was reduced to less than 10 kΩ. The preprocessing was performed offline in MATLAB (2019b) using the Fieldtrip toolbox (v. 20230318, Oostenveld et al., 2011). All data were down-sampled to 250 Hz, referenced to the average of all electrodes, and band-pass filtered between 0.5 to 40 Hz. Artifacts (blinks, eye movements and muscle contractions) were removed by visual inspection and by independent component analysis (FASTICA; Hyvärinen and Oja, 2000). Continuous data were segmented into two main distinct epochs. The first, called “decoding epoch” was time-locked to S1 offset and sorted according to *stimulus type* and *S1 duration*. The contrast was not considered as “sorting” feature since it showed no significant effect on behavioral data (see “4.1 Behavioural results”). The ‘decoding’ epoch spanned from S1 offset to 0.5 s after S1 offset (before reproduction onset) and was baseline-corrected to the 0.1 s around S1 offset (-0.05 to 0.05 s). The second, ‘encoding’ epoch comprised the last 0.2 s of S1 presentation and was baseline corrected using the 0.2 s preceding S1 onset. All epochs were further screened for residual artifacts via visual inspection, leading to the rejection of 156 trials (on average 1.65% of trials per participant). The EEG analyses focused on both encoding and decoding epochs, with greater emphasis on the decoding epoch for several reasons: (a) it is the phase of the task in which the different stimulus types are physically identical; (b) it is when the stimulus duration becomes available to the observer; (c) unlike the reproduction phase, it is free from concurrent processes such as memory retrieval or motor preparation; and (d) it is the phase on which previous studies have primarily focused (Ofir & Landau, 2022).

#### 3.3.3 Event related potential (ERP) analysis

Event-related potentials (ERPs) were computed separately for *stimulus type* and *S1 duration*. To explore the interaction between stimulus type and duration, we conducted a General Linear Model (GLM) analysis using EEGLAB (Delorme & Makeig, 2004) in conjunction with the LIMO toolbox (Pernet et al., 2020). In this analysis, the combination of *stimulus type* and *S1 duration* was used as categorical predictors, resulting in a total of nine predictors (i.e., 3 stimulus types by 3 S1 durations). At the first level of analysis, beta values for each predictor were estimated in each individual subject. At the second, group-level analysis, we performed a two-way Analysis of Variance (ANOVA) on the nine betas estimated for each of the twenty-one participants. The statistical significance was assessed using spatial-temporal clustering, following the pipeline proposed by Pernet et al. (2021).

#### 3.3.4 Representational Similarity Analysis

Representational similarity analysis (RSA, Kriegeskorte et al., 2008) was used to identify the EEG correlates of the filled-duration illusion measured in behavior. First, for each participant we constructed a behavioral Representational Dissimilarity Matrix (RDM). In this 9×9 behavioral-RDM, each element represented the dissimilarity - quantified as Euclidean distance - between the mean reproduction performance (i.e., the bias - see “3.3.1 Behavioral data analysis”) for each pair of conditions (a combination of *S1 duration* and *stimulus type*). Specifically, mean bias was computed as the average across trials, separately for each combination. Only trials that were not excluded by epoch rejection were considered. Similarly, for each participant, we generated EEG-derived RDMs, one for each electrode and each time point (for a total of 7750 EEG-RDMs for each participant). Each element of these EEG-RDMs represented the dissimilarity between the EEG responses of pairs of conditions. This difference was computed on trial grand-averages. Furthermore, we generated two theoretical RDMs that matched the dimensions of the empirical RDMs. The first, referred to as the stimulus-RDM, was designed to express the dissimilarity between stimuli with arbitrary values. The second, referred to as the physical duration-RDM, was constructed by using differences in S1 durations between pairs. Notably, in this case the values reflect the behavior of an ideal observer who would reproduce duration without any bias, behaving as a “perfect clock”. For subsequent analysis, we focused exclusively on specific sub-quadrants of the RDMs, as highlighted in Figure 6A. These sub-quadrants specifically represented the contrast between the empty and the filled and flanked stimulus conditions, as this comparison yielded a significant effect in the behavioral analysis (see Results section). At each time point and each electrode, we calculated the Spearman correlation between the EEG-derived RDMs and the corresponding elements in the three other matrices i.e., the behavioral-RDM, the stimulus-RDM and the duration-RDM. Thus, for each participant, we obtained three matrices (N_electrodes_ x N_time points_) containing correlation values, which were then Fisher-transformed to z-scores. The z-transformed correlation values were then tested against zero with a cluster-based one-sample T-test. Finally, the correlation scores between EEG RDMs and behavioral RDMs were contrasted with with those between EEG RDMs and physical duration RDMs using a cluster-based permutation T-test. This constituted our main test of interest, as it identifies where the perceptual bias observed behaviorally is reflected in EEG activity. Specifically, we examined whether EEG signals encode perceptual distortions induced by stimulus type *in addition to* the representation of physical duration. The entire RSA pipeline is schematically illustrated in Figure 6.

## 4 Results

### 4.1 Behavioural results

In this study, we investigated how stimulus format (or sensory load) influences perceived duration. Participants performed a duration reproduction task under three distinct stimulus conditions characterized by a different sensory load (i.e., filled, flanked and empty temporal intervals) and we measured their reproduced durations.

Reproduction performance (quantified as bias, Figure 1B) was analysed with a LME model with *S1 duration, stimulus type* and *contrast* as fixed predictors and *participant* as a random intercept factor (formula: *bias* ∼ *stimulustype* ∗ *duration* ∗ *contrast* + (1|*subjectID*)). A type III ANOVA on model estimates (marginal *R*2 = 0.57, conditional *R*2 = 0.74) revealed a main effect of *stimulus type* (*F* (2, 340) = 69, 24, *p <* .0001) and a main effect of *S1 duration* (*F* (2, 340) = 345.97, *p <* .0001). Post-hoc pairwise comparisons (Tukey-adjusted) on the estimated marginal means for each stimulus type, revealed that the reproduction bias differed between the empty and the filled interval (empty-filled: *t*(340) = −8.78, *p <* .0001) and between the empty and flanked interval (empty - flanked: *t*(340) = −11.17, *p <* .0001). The difference between the filled and flanked intervals also reached significance, although with much smaller statistical significance (filled-flanked: *t*(340) = −2.39, *p* = .05).

Next, we examined the coefficient of variation (CV) as a measure of variability in reproduction performance (Figure 1C). We fitted a LME model with formula *CV* ∼ *stimulustype* ∗ *duration* ∗ *contrast* + (1|*subjectID*). Type III ANOVA on model estimates (marginal *R*2 = 0.14, conditional *R*2 = 0.47) showed that CV was significantly affected by both *stimulus type* (*F* (2, 340) = 24.48, *p <* .001) and *duration of S1* (*F* (2, 340) = 22.26, *p <* .0001), as shown in Figure 1C. Post-hoc tests (Tukey-adjusted) on the estimated marginal means for each stimulus type revealed significant differences in CV between empty and filled intervals (empty - filled: *t*(340) = 5.67, *p <* .0001) and between empty and flanked (empty - flanked: *t*(340) = 6.39, *p <* .0001). The contrast between filled and flanked intervals was not significant (filled - flanked: *t*(340) = 0.724, *p* = .75).

Stimulus Contrast did not significantly affect performance, neither in bias nor variability. The main effect of contrast and its interactions with stimulus type and S1 duration were not significant. Accordingly, contrast was excluded from subsequent analyses to optimize statistical power..

### 4.2 ERP results

#### Decoding epoch

This analysis focused on the temporal window immediately following the offset of the template stimulus (S1). We refer to this as the ‘decoding’ epoch, as this is the moment when the duration of the presented interval has elapsed and should be read out. This window of interest spanned from S1 offset up to 0.5 s later, before reproduction onset. The reason for choosing this specific window was two-fold: first, this was the moment when stimulus duration became available to the observer, and unlike the reproduction phase, it was free of concurrent processes such as memory retrieval or motor preparation. Second, at stimulus offset, all three experimental conditions (i.e., the different stimulus types) were physically identical, since nothing was projected on screen before the reproduction onset. To explore the effects of *stimulus type* and *S1 duration*, for each subject, we conducted a General Linear Model (GLM) on the EEG signal during the decoding epoch (from S1 offset to 0.5 s after).

The group-level ANOVA on the nine GLM estimated predictors (i.e., the combination of 3 S1 durations and the 3 stimulus types) resulted in a main effect of *S1 duration* (see Figure 2A, left panel). This effect was driven by a large cluster of electrodes extending in space from the occipital to the frontal regions and in time from 0.26 to 0.5 s after stimulus offset (maximum *F* (2, 19) = 70.46, *p* = 0.001, cluster-corrected), and whose peak was 0.285 s in channel P2. From the visual inspection of the topography (2B, left panel), it seemed that *S1 duration* was associated with an increase in the effect in both occipital and frontocentral electrodes, most prominently in a temporal window spanning from 0.1 to 0.5 s. To better qualify the effect of *S1 duration* we inspected the average ERPs in two subsets of electrodes: a frontocentral set (F1, Fz, F2, FC1, FCz, FC2, Cz,C1, C2), that aligned with prior findings in the literature (Bueno & Cravo, 2021; Ofir & Landau, 2022); and an occipital set (Oz, POz, O1, O2, PO3, PO4, PO7, PO8), which was linked to the specific demands of the visual task. As shown in Figure 3A, *S1 duration* modulated ERP amplitude in similar temporal windows and in a similar fashion across all stimulus conditions. Interestingly, the direction of this modulation was reversed in occipital compared to frontal electrodes: in occipital electrodes, shorter S1 durations were associated with more negative ERP amplitudes, whereas in frontocentral electrodes, the reverse was true (shorter S1 durations led to more positive amplitudes, Figure 3A).

**Figure 2:**
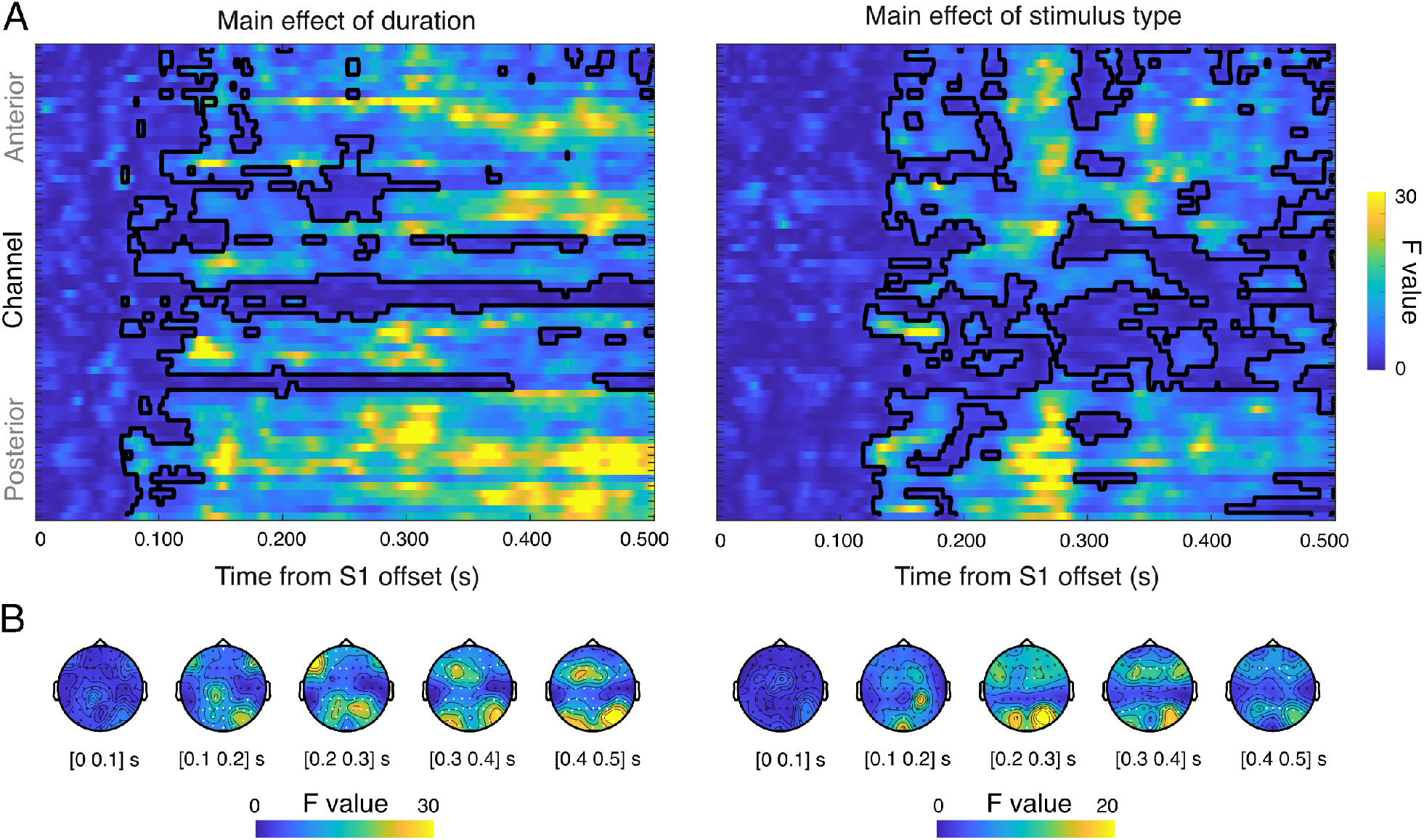
Results of the ANOVA ran on GLM estimates for the decoding epoch. **A)** F-score for the effect of *S1 duration* (left) and *stimulus type* (right) in the decoding epoch (i.e. 0.5 s following S1 offset) for each channel and time point. Black contours indicate points of statistical significance. **B)** Topographies of the effects are reported in windows of 0.1 s; highlighted electrodes yielded a significant effect throughout the 0.1 s window.

**Figure 3:**
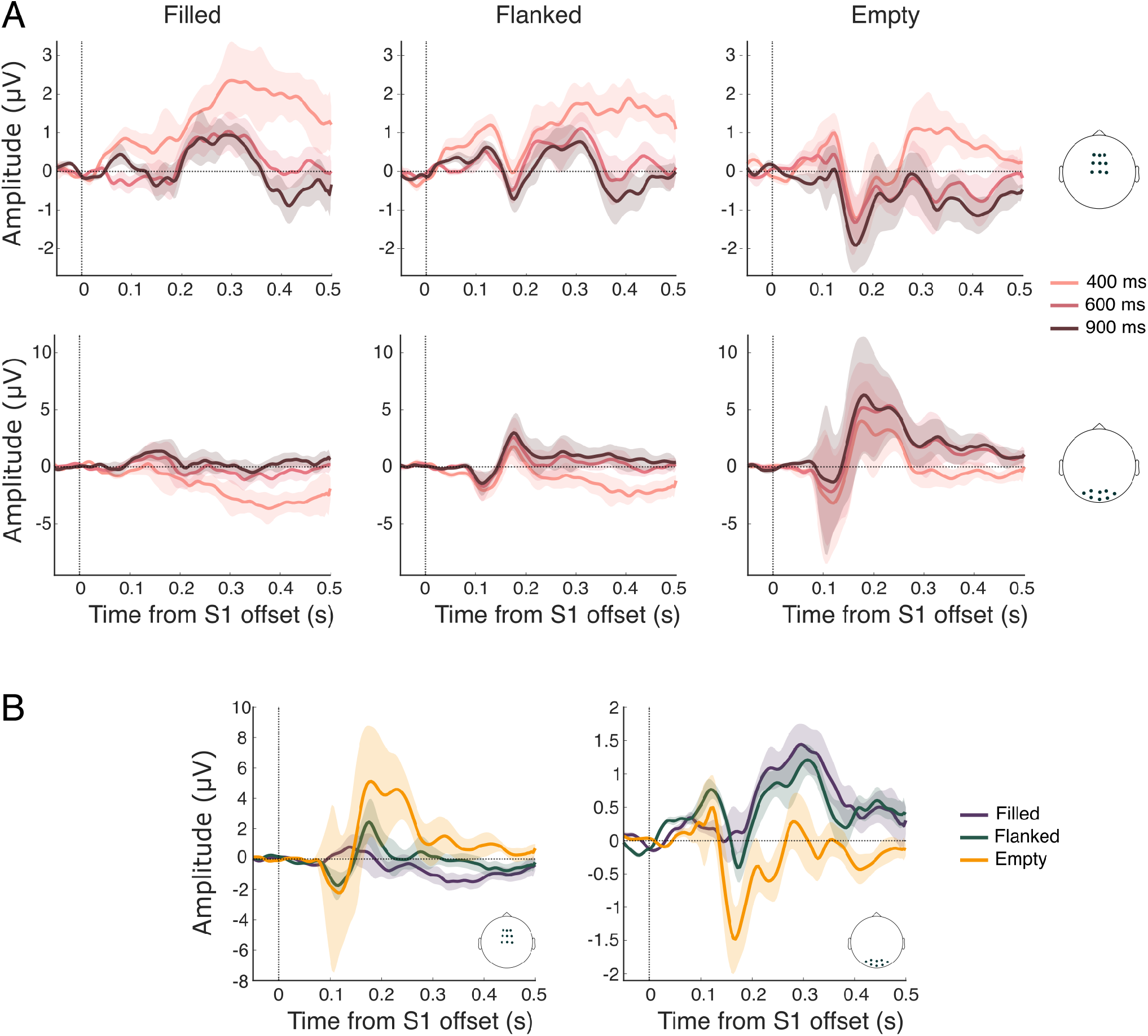
Grand average ERP waveforms aligned to S1 offset (decoding epoch). A) Waveforms are divided by S1 duration and displayed for frontocentral (top row) and occipital (bottom row) electrodes, across stimulus conditions (Filled, Flanked and Empty). B) Waveforms are displayed by electrodes subsets (occipital, left; frontocentral, right), pooled across S1 durations and divided by stimulus type. In both panels, shaded areas indicate the Standard Error of the Mean.

The ANOVA on the GLM estimated parameters also showed a main effect of *stimulus type* (see Figure 2A, right panel). The first significant time point was 0.08 s after stimulus offset whereas last one was 0.499 s after stimulus offset. The peak (*F* (2, 19) = 66.52, *p* = 0.001) was at 0.131 s in channel CP4. The visual inspection of the topography shows that in a temporal window between 0.200 and 0.300 s after S1 offset, stimulus type leads to a greater effect in occipital electrodes (see Figure 2B, right panel). When average ERPs in frontocentral and occipital sets were grouped by *stimulus type*, collapsing across durations, a modulation according to the sensory load of the temporal intervals emerged (Figure 3B): the amplitude of the ERPs associated with filled stimuli was more similar to flanked than empty.

The interaction *stimulus type* × *S1 duration* did not reach statistical significance.

#### Encoding epoch

We also analyzed the data referring to the stimulus presentation epoch (the ‘encoding epoch’), to explore the possibility that differences in temporal processing between stimulus types would already emerge during S1 presentation. However, to avoid the confounding effect of having strong ERPs responses at stimulus onset and also to equate the temporal window of interests for different S1 durations, we chose to look at the last 0.2 s before interval offset.

The group level results from the ANOVA performed on individual GLM beta estimates showed significant effects of the two predictors (*stimulus type* and *S1 duration*) and their interaction (Figure 4). The analysis identified a main effect of *S1 duration* throughout the last 0.2 s of the interval, with a peak at electrode F5, 0.168 s before the stimulus offset (*F* (2, 60) = 62.41, *p* = .001). A main effect of *stimulus type* was also identified throughout the analyzed window, peaking 0.016 s before stimulus offset at electrode P4 (*F* (2, 60) = 53.51, *p* = 0.001). Finally, an interaction effect emerged between 0.2 and 0.1 s before stimulus offset, with a peak at electrode P7, 0.192 s before S1 offset (*F* (4, 17) = 20.97, *p* = 0.025). The topographies show that, whereas stimulus duration affected the entire epoch, the effect of stimulus type was greater in the last 0.1 s before stimulus offset in parieto-occipital and frontocentral electrodes (Figure 4B).

**Figure 4:**
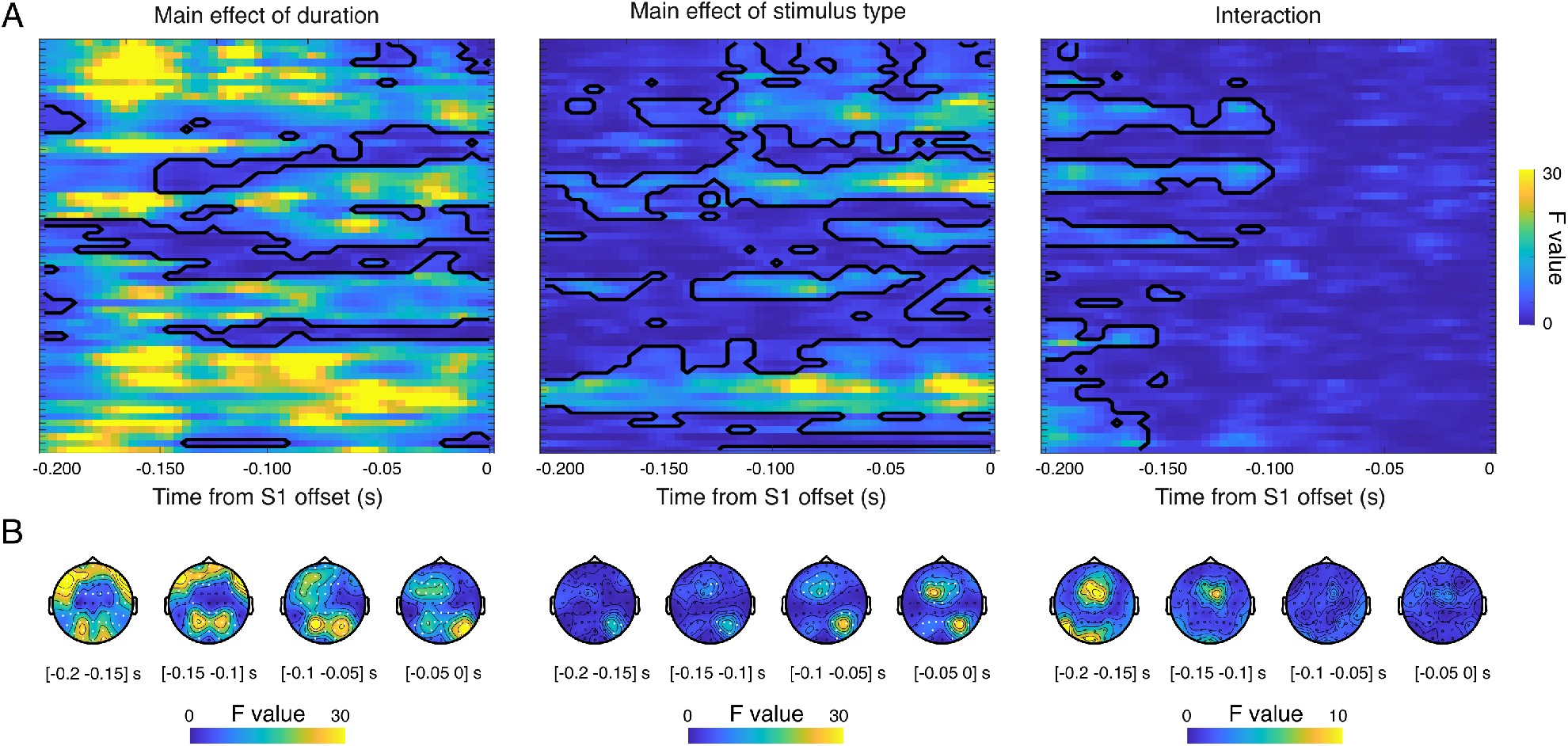
Results of the ANOVA ran on GLM estimates for the encoding epoch. **A)** F-score for the effect of *S1 duration* (left), *stimulus type* (middle) and their interaction (right) in the encoding epoch (i.e. the last 0.2 s of S1 presentation). Results are reported for all time points and all channels. Black contours indicate points of statistical significance. **B)** Topographies of the effects are reported in windows of 0.1 s; highlighted electrodes yielded a significant effect throughout the 0.1 s window.

For completeness, we also looked at the ERPs time-locked to S1 onset and throughout the stimulus presentation window, averaged among the same frontocentral and occipital sets of electrode as for the decoding. Figure 5 displays average ERPs for different S1 durations (separate panels) and stimulus types (color coded lines) recorded in the same subset of electrodes used for the decoding window. Surprisingly, the average ERPs were very similar across durations and stimulus types. Specifically, these deflections were ordered according to the physical/sensory load difference between the stimulus types. The average ERPs for filled intervals were more similar to those of the flanked compared to the empty stimuli, and the ERPs of the empty intervals were more similar to those of the flanked compared to filled intervals. As in the decoding epoch, the sign of these modulations was reversed in frontocentral compared to occipital electrodes. Overall, the ERP data recorded in both the encoding and decoding stages showed a modulation of the signal amplitude associated to both *S1 duration* and *stimulus type*.

**Figure 5:**
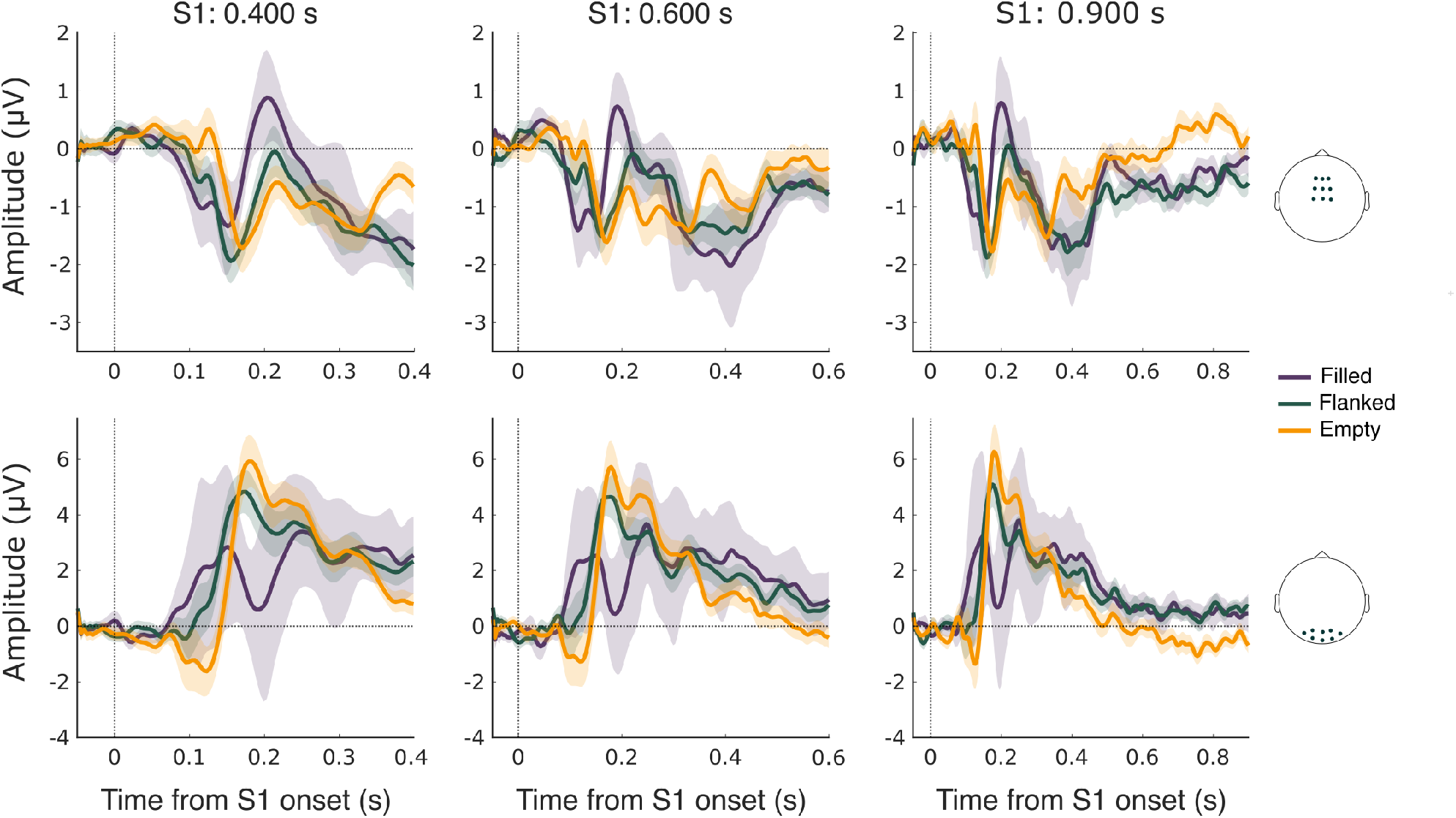
Grand average ERP waveforms aligned to the S1 onset (encoding epoch). ERPs are reported for frontocentral (top row) and occipital (bottom row) electrodes. Stimulus condition is color-coded and plots are divided by S1 duration. Shaded areas indicate the Standard Error of the Mean.

### 4.3 RSA results

The following step aimed to explore the relationship between behavior and EEG data, specifically seeking correlates of *perceived* rather than physical duration. Since different stimulus types led to differences in performance, especially when comparing filled and flanked versus empty stimuli, we employed Representational Similarity Analysis to identify EEG signature of these performance differences. The first step of this analysis required the computation for each subject of 4 distinct Representational Dissimilarity Matrices (RDM), two empirical and two theoretical (Figure 6A). The empirical matrices were a behavioral RDM and an EEG-derived RDM. In these 9×9 RDMs, each element represented the dissimilarity - quantified as Euclidean distance - between either the reproduction performance (bias - see “3.3.1 Behavioral data analysis”) or the EEG signal for each pair of conditions (a combination of S1 duration and stimulus type). For each subject we had one behavioral-RDM and an EEG-RDM for each electrode and each time point. The two theoretical RDMs, that matched the dimensions of the empirical ones, were the stimulus type and the physical duration -RDMs. The first (stimulus-RDM) was designed to express the dissimilarity between filled, flanked and empty intervals with arbitrary values. The second (duration-RDM), was built by using differences in S1 durations between pairs as if an ideal observer would reproduce them without any bias. We next computed, for each time point and each electrode, Spearman correlations between the EEG-RDMs and each of the three other matrices: the behavior-RDM, the stimulus-RDM and the duration-RDM. By correlating EEG RDMs to both empirical and theoretical RDMs, we assessed the extent to which these representational geometries overlapped. Thus, for each participant, we obtained three matrices (N_electrodes_ x N_time points_, Figure 6B) containing correlation values, which were then Fisher-transformed to z-scores. It is important to note here that these correlations were performed exclusively in specific sub-quadrants of the RDMs, as shown in Figure 6A. These were indeed the sub-quadrants representing the contrasts where we observed the greater difference in performance i.e., between empty and filled conditions and empty and flanked conditions (see “4.1 Behavioural results”). It is important to emphasize here that the representational geometries of behavior and of pure duration are necessarily very similar, since duration was the main factor influencing behavior given the task at hand. To determine whether the EEG signal reflected perceived duration (indexed by behavior) *more* than physical duration, we tested the difference between these two correlation matrices. The results of this test are shown in Figure 6C. The same pipeline was applied to both the decoding and the encoding windows.

**Figure 6:**
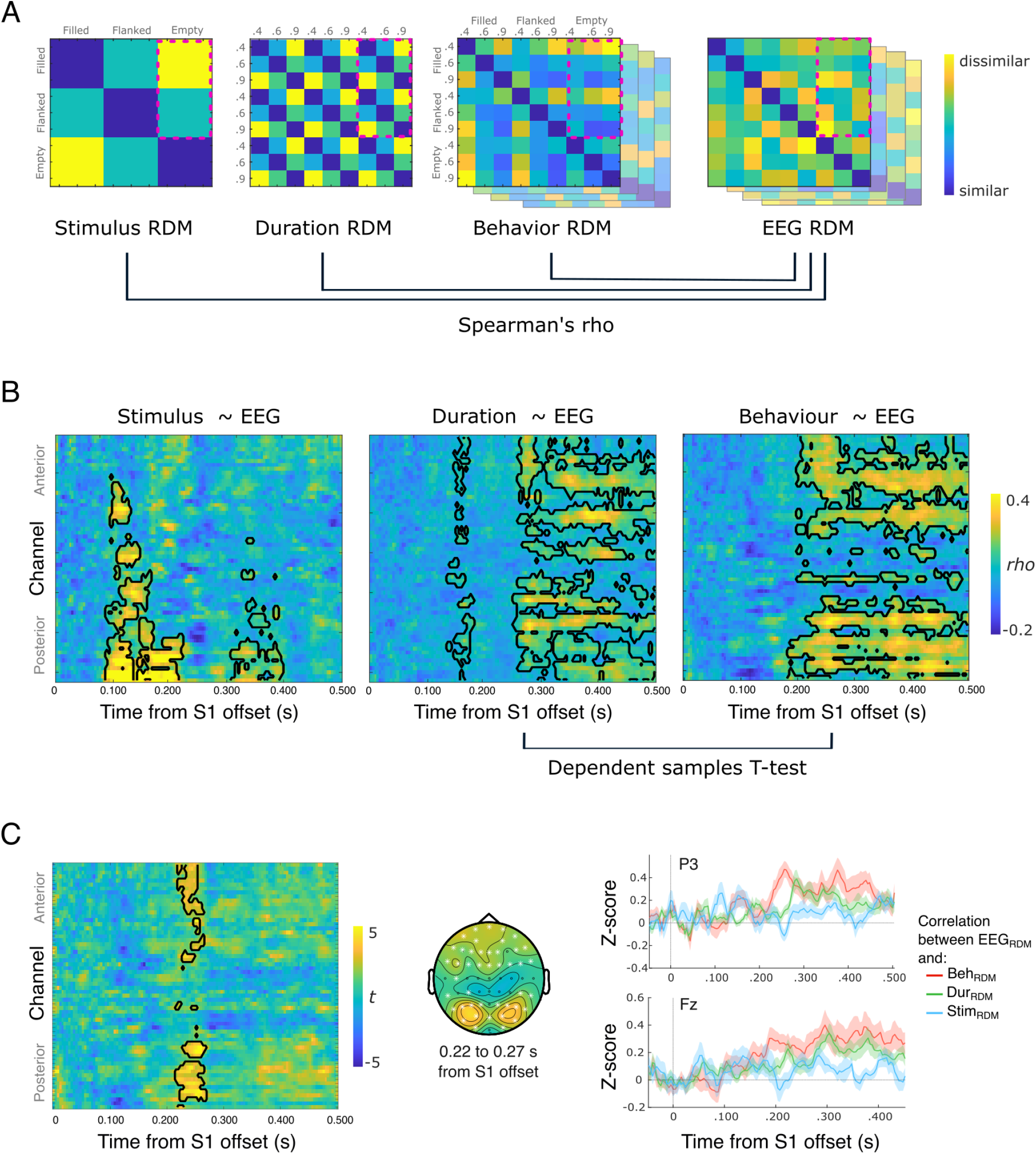
Pipeline and results from RSA on the decoding epoch. **A)** Representational Dissimilarity Matrices (RDMs) illustrating the representational geometry of theoretical matrices based on stimulus type and physical duration information, juxtaposed with the representational geometry derived from behavioral performance and EEG data. Spearman’s rho correlations were calculated between the EEG RDM and each of the following: stimulus RDM, duration RDM, and behavior RDM. The resulting rho values were Fisher-transformed, and the corresponding z-scores were tested against zero (see panel B). **B)** Results from the correlations between stimulus-RDMs and EEG-RDM (left panel), duration-RDM and EEG-RDM (middle panel) and behavior-RDM and EEG-RDM (right panel). For each time point and electrode, we report the correlation coefficient. Z-transformed correlation coefficients were tested against zero with a cluster based dependent samples T-test. Black contours indicate the channels and time points where the correlation was significant (p¡.05, cluster corrected). **C)** Contrast (computed as cluster-based dependent samples T-test) between the correlation matrix representing similarity between EEG-RDMs and behavior-RDM and the correlation matrix representing the similarity between EEG-RDMs and duration-RDM. In the left panel, t-scores are displayed for each channel and time point; contours indicate the significant cluster, centered at .250 s after offset and localized in parietal and occipito-parietal electrodes. Here, the similarity between EEG and behavior is higher than the similarity between EEG and duration. The middle panel displays the topographic distribution of the average t-value within the significant cluster; white stars indicate electrodes that fall into the cluster. In the right panel, for visualization purposes, we report similarity values (z-transformed Spearman’s rho) between EEG-RDMs and each of the three matrices above at two example electrodes (PO3 and P6). Lines indicate the group average of z-value and shaded areas indicate the Standard Error of the Mean.

#### Decoding epoch

First, we tested the significance of correlation scores at group level. Correlation values between the stimulus-RDM and the EEG-RDMs were significantly above zero in two clusters. The first cluster (*p <* 0.001) spanned from 0.09 to 0.22 s after S1 offset, with a peak at electrode O2 (*rho* = 0.70 at 0.11 s after S1 offset) and an average *rho* = 0.25 within the cluster. The second group (*p <* 0.001) ranged from 0.31 to 0.4 s after S1 offset, with a peak at electrode F3 (*rho* = 0.28) 0.33 s after S1 offset and an average *rho* = 0.17 within the cluster. Correlations between the duration RDM and EEG RDMs were also statistically significant in two spatiotemporal clusters. The first cluster (*p <* 0.001) spanned 0.25 to 0.5 s after S1 offset, with an average *rho* = 0.16 and a peak at electrode FC1 (*rho* = 0.34), at 0.38 s. The second cluster (*p <* 0.05) spanned 0.14 to 0.18 s after S1 offset, peaking at electrode C5, 0.16 s after S1 offset (*rho* = 0.21), with an average *rho* = 0.11. Finally, correlation values between the behavior-RDM and the EEG-RDMs were significant within a cluster spanning 0.18 to 0.5 s after S1 offset, with an average *rho* = 0.22 and a peak in correlation at electrode P1 (*rho* = 0.41) 0.43 s after S1 offset.

By visually inspecting the time-course of similarity values at example electrodes (Figure 6C, right panel), it is evident that stimulus type was strongly represented at early latencies in occipital electrodes, whereas representations of pure duration and behavior emerged at later latencies.

Then, we investigated whether the correlation values between EEG-RDMs and behavior-RDMs were higher than those between EEG-RDMs and duration-RDMs, using a cluster-based dependent samples T-test. We found a significant difference driven by two positive clusters. The first cluster (*p* = 0.008) spanned from 0.22 to 0.27 s after S1 offset, and included parietal and occipito-parietal electrodes (6C, left and middle panels); the average effect size within the cluster, measured by Cohen’s *d*, was *d* = 1.01, while the maximal effect was found at 0.26 s at electrode P3 (*d* = 1.37). The second cluster (*p* = 0.04) spanned from 0.22 to 0.26 s after S1 offset, encompassing frontal and frontocentral electrodes; the average effects size within the cluster was *d* = 0.87, and the peak was found at 0.26 s at electrode F6 (*d* = 1.34). These results indicate that, in this time window and at posterior electrodes, the EEG signal reflected perceived rather than physical duration, as the correlation of EEG-RDMs with behavior-RDMs was significantly stronger than the correlation with duration-RDMs.

#### Encoding epoch

The main results of the RSA analysis applied to the last 0.2 s of the stimulus presentation epoch are shown in Figure 7. Significant correlations between EEG-RDMs and stimulus-RDMs were observed in two small clusters (Figure 7A, left panel). The first cluster included anterior and fronto-parietal electrodes, and extended from 0.16 to 0.14 s before S1 offset, peaking at electrode FP1 (*rho* = 0.25), 0.14 s before S1 offset, with an average *rho* = 0.16 within the cluster. The second cluster peaked at electrode F6 (*rho* = 0.22), 0.06 s before S1, with a mean *rho* = 0.14 across the cluster. The duration-RDM correlated significantly with the EEG-RDMs across all electrodes and throughout the analyzed window (starting 0.2 s before S1 offset, Figure 7A, middle panel). The peak in correlation was found at electrode CP5 (*rho* = 0.50), 0.16 s before S1 offset and the average correlation coefficient within the cluster was *rho* = 0.21. Similarly, behavior-RDMs correlated significantly with EEG-RDMs across all electrodes and throughout the entire analyzed window (Figure 7A, right panel), with an average correlation coefficient *rho* = 0.23 within the cluster and a correlation peak at electrode F7 (*rho* = 0.45), 0.11 s before S1 offset.

**Figure 7:**
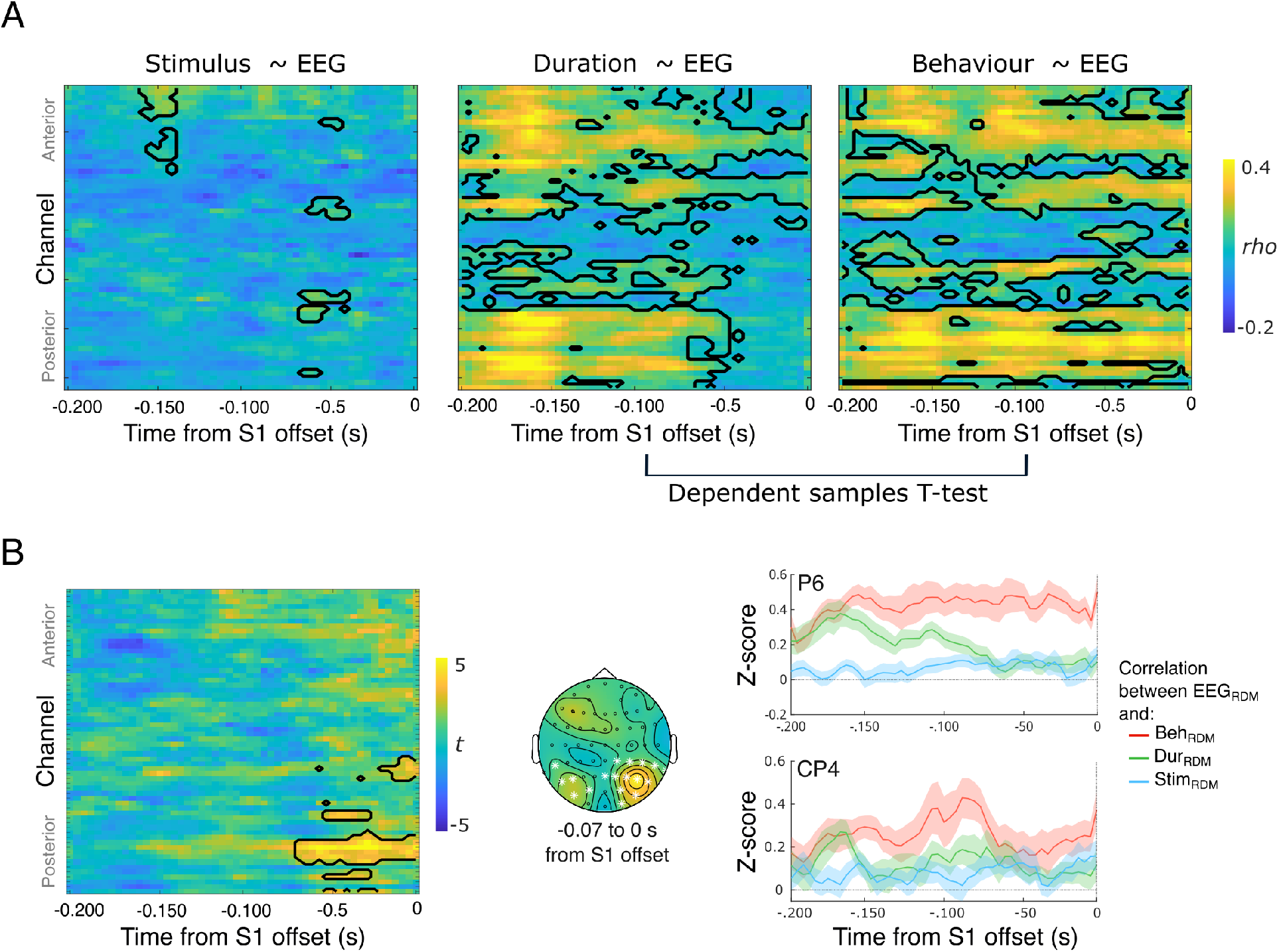
Results from RSA on the encoding epoch. **A)** Results from the correlations between stimulus-RDMs and EEG-RDMs (left panel), duration-RDM and EEG-RDMs (middle panel) and behavior-RDMs and EEG-RDMs (right panel).For each time point and electrode, we report the correlation coefficient. Z-transformed correlation coefficients were tested against zero with a cluster based dependent samples T-test. Black contours indicate the channels and time points where the correlation was significant (p¡.05, cluster corrected). **B)** Results of the cluster-based permutation test on correlation scores between EEG-RDMS and duration-RDM, and EEG-RDMS and behavior-RDMs. The left panel displays t-scores in space (electrodes) and time (seconds); the significant cluster is represented by black contours. The middle panel displays the topographic distribution of the average t-value within the cluster; white stars indicate electrodes that fall into the cluster. In the right panel, two example electrodes (P6 and CP4) showing the time-course of similarity (expressed as z-transformed correlation coefficients) between EEG-RDMs and both theoretical and behavioral RDMs are displayed.

Finally, we examined whether the EEG signal represented behavior more strongly than physical duration, similar to the analysis performed for the decoding epoch. Results showed that correlation values between EEG-RDMs and behavior-RDMs were significantly higher than correlation values between EEG-RDMs and the duration-RDM in the last 0.07 s of the stimulus presentation epoch (Figure 7B, left panel), at parietal and occipital electrodes (Figure 7B, middle panel). The peak of the effect was found 0.03 s before stimulus offset at electrode P6 (*d* = 1.35); the average effect size over the largest cluster was *d* = 0.79.

## 5 Discussion

In the present study, we sought to explore the neural differences in the duration processing of stimuli with different sensory load and to identify the neural correlates of the perceptual biases associated with these different stimulus configurations. To this end, participants were asked to reproduce visual temporal intervals of three different stimulus types - filled, flanked and empty intervals - while EEG was recorded.

As expected, our behavioral findings revealed differences in reproduced duration between stimulus conditions, aligning with the ‘filled duration illusion’ phenomenon (Bratzke et al., 2017; Thomas & Brown, 1974; Wearden et al., 2007). Specifically, participants consistently under-reproduced empty intervals compared to both filled and flanked intervals. Interestingly, a similar behavioral performance was observed for filled and flanked intervals compared to empty intervals. This result suggests that, despite the presence of highly salient markers at onset and offset, the amount of sensory input provided *throughout* the stimulus is crucial to shape perception. Behavioral differences between the empty condition and the other two stimulus configurations were not only measured in mean reproduced duration, but in reproduction variability as well (as expressed by the Coefficient of Variation). Specifically, empty intervals were associated to higher variability, in line with effects previously reported in the auditory modality (Rammsayer, 2010; Rammsayer & Lima, 1991).

At the EEG level, we characterized the signature of temporal processing across stimulus conditions. Event-related potentials (ERPs) following S1 offset exhibited neural responses that varied systematically with stimulus duration. Namely, interval duration modulated brain responses in a gradual and ordered fashion under all three stimulus conditions, most prominently in the occipito-parietal region. Previous studies reported similar duration effects in electrophysiological activity after stimulus offset (Baykan et al., 2023; Bueno & Cravo, 2021; Kononowicz & van Rijn, 2014; Ofir & Landau, 2022; Özoğlu & Thomaschke, 2023). However, these studies identified duration effects in a decisional phase, either because single interval tasks were used (Baykan et al., 2023; Ofir & Landau, 2022; Özoğlu & Thomaschke, 2023) - and thus temporal encoding could not be dissociated from temporal decisions by design - or because the reported effects refer to the second interval of the pair (Bueno & Cravo, 2021; Kononowicz & van Rijn, 2014). Instead, our analysis focused on epochs that were devoid of decision-making processes. When presented with the template stimulus (S1), participants were not comparing it to a previously presented stimulus or reference, but solely processing its duration for the purpose of reproduction. Duration effects on ERPs at stimulus offset in the context of a reproduction task have been reported in the auditory modality. Damsma et al. (2021) report ERP modulations by duration at central electrodes. However, their results differ from ours in a key respect, as the relationship between duration and voltage is reversed: in our data, shorter durations corresponded to higher (more positive) signal amplitudes at frontocentral electrodes, whereas the opposite pattern was observed in their study. This discrepancy could be due to a different subset of analysed electrodes, but it may also reflect modality-specific differences in the topography of time-related effects in the auditory versus visual domains. Our results align most closely with recent findings by Bueno et al. (2024), who showed that interval duration could be successfully decoded from EEG activity both at the end of stimulus presentation and after its offset. Notably, their ERPs at S1 offset closely resemble our findings, further supporting the idea that neural activity at this stage reflects temporal processing.

While Bueno et al. (2024) primarily focused on generalizing duration effects across timing tasks (discrimination and reproduction), our study extends this finding to different stimulus configurations. We found that the stimulus’ sensory load significantly affected the EEG signal both during and after stimulus presentation. However, we argue that these effects do not imply distinct timing mechanisms. In the post-offset temporal window (the ‘decoding epoch’), we found that duration is represented in the same way across stimulus conditions. The absence of an interaction effect between stimulus type and duration following S1 offset suggests a consistent, unified process for duration representation, that is not significantly altered by the specific sensory configuration of the interval. With respect to the stimulus presentation epoch (the ‘encoding epoch’), and consistent with previous research (Pfeuty et al., 2008), we found voltage differences between stimulus conditions. This result emphasizes the brain’s sensitivity to varying stimulus configurations, in the absence of concurrent comparison or decisional processes. The observed voltage differences likely reflect the physical differences between stimuli and lead to differences in both the encoding and decoding of time.

Our findings support the importance of sensory load in defining the duration percept. By relating EEG activity patterns to behavioral patterns with RSA, our findings support the idea that electrophysiological differences induced by the specific physicality of stimuli shape the forthcoming behavioral performance - which is a proxy of perceived duration. Specifically, at early post-offset latencies, neural patterns predominantly reflected stimulus type; as the representation of stimulus type decayed, the neural representation shifted to reflect physical and perceived duration. Crucially, it was at the intersection of these representations that behavioral distortions were represented *beyond* pure duration, most prominently at occipito-parietal electrodes. Furthermore, we found correlates of perceived duration already before the end of the interval presentation, in an epoch (the last 0.1 s) where effects of stimulus type on signal were found, indicating that this early signal shapes the forthcoming behavioral performance. Overall, these results highlight the integration of stimulus-induced biases in the neural representations of time and indicate that distortions in time perception do not originate from late decision making or memory processes, but rather are inextricably related to the way in which sensory information is encoded and represented-that is, an early and sensory stage of temporal processing.

To sum up, our findings align with previous research showing that the stimulus sensory load significantly influences temporal perception (Reinartz et al., 2024; Tonoyan et al., 2022). However, the EEG correlates of duration processing were consistent across stimulus configurations, suggesting the existence of a shared duration processing mechanism for filled, flanked and empty visual stimuli. Furthermore, we show that signatures of temporal processing reflect perceptual biases induced by different stimulus configurations. Rather than positing that differences in perceived duration arise from differences in timing strategies, we interpret these biases in a more parsimonious way, namely that they originate from the integration of stimulus-specific sensory information with timing processes. We conclude that stimulus-induced biases are incorporated into temporal representations at an early stage, emphasizing the dynamic interplay between sensory information and the brain’s encoding of the temporal dimension.

## Acknowledgements

D.B. acknowledges financial support from the European Research Council–ERC (Grant Agreement No 682117, BiT-ERC-2015-CoG), the Italian Ministry of University and Research under the call PRIN2022 (project ID: CCPJ3J).

